# Phase-specific stimulation of the human brain with real-time measurement instead of prediction

**DOI:** 10.1101/2023.04.20.537612

**Authors:** Robert Guggenberger, Julian-Samuel Gebühr, Marius Keute, Alireza Gharabaghi

**Author notes:** Correspondence Institute for Neuromodulation and Neurotechnology, University Hospital and University of Tuebingen, Otfried-Mueller-Str.45, 72076 Tuebingen, Germany. Email address.

## Abstract

**Background:** The responsiveness of the human brain to external input fluctuates. Timing the external perturbation with regard to the oscillatory brain state may improve the intended stimulation effects. However, current brain state-dependent interventions targeting phases of the oscillatory cycle need to apply prediction algorithms to compensate for latencies between measurement and stimulation, and are therefore imprecise.

**Objective:** We investigated the phase-specific precision of a novel non-predictive approach on the basis of integrated real-time measurement and brain stimulation.

**Methods:** Applying a simulation, we estimated the circular standard deviation (SD) to hit 2, 4, 8, 16 or 32 equidistant phase bins of the oscillatory cycle with high precision. Furthermore, we used electroencephalography-triggered transcranial magnetic stimulation in healthy subjects to empirically determine the precision of hitting the targeted phase of the oscillatory cycle for 10 different frequencies from 4Hz to 40Hz using our approach.

**Results:** The simulation revealed that SDs of less than 17.6°, 9.7°, 5.1°, 2.5°, and 1.3° were necessary to precisely hit 2, 4, 8, 16, and 32 distinct phase bins of the oscillatory cycle. By completing measurement, signal-processing and stimulation with a round-time of 1ms, our empirical approach achieved SDs of 0.4° at 4Hz to 4.3° at 40Hz. This facilitates selective targeting of 32 phases (at 4Hz), 16 phases (at 8, 12, 16, 20, 24Hz) and 8 phases (at 28, 32, 36, 40Hz), respectively.

**Conclusion:** Integrated real-time measurement and stimulation circumvents the need for prediction and results in more precise phase-specific brain stimulation than with state-of-the-art procedures.

## Introduction

The growing interest in brain state-informed interventions in neuroscience and therapy is driven by the motivation to achieve more predictable stimulation effects and neuroplastic changes. Specifically, EEG-triggered transcranial magnetic stimulation (TMS) has been applied to repetitively target sensorimotor rhythmic activity or oscillatory up- and down phases in order to induce increased corticospinal excitability in the human motor cortex, albeit with contradictory findings.^1-3^

To identify the impact of stimulation timing on fluctuations of motor-evoked potential (MEP) amplitudes, the pre-TMS power and phase at the site of stimulation have been investigated with post-hoc analysis.^4^ Specifically, a 40-70% MEP amplitude increase was detected across different beta power levels, thereby implying a certain robustness against imprecise stimulation timing. By contrast, the phase-modulation was critically dependent on precisely timing the stimuli to specific phases of the oscillatory cycle. Importantly, when this phase specificity was achieved, a MEP increase of 180% could be attained.^4^

Different methods are currently being applied to predict the oscillatory phase for state-informed stimulation: Autoregressive (AR) model-based approaches;^5^ Kalman filter-based AR approaches for long-term predictions (> 100 ms);^6^ least mean square (LMS)-based AR approaches for adaptive predictions with recurrent updates;^7^ approaches that utilize pre-learned features;^8^ and machine learning-based approaches on the basis of training data.^9^ Importantly, these approaches often apply their phase predictions to lower frequency oscillations, since these have inherently less strict demand for temporal precision than higher frequency oscillations. The higher the frequency, the more challenging the task of targeting a specific phase bin, since the round-trip time needs to be less than the width of the bin for the target frequency. This challenge becomes particularly apparent when more precise phase targeting than hitting two opposite phases (e.g., peak vs. trough) in the alpha band is necessary to achieve the intended stimulation effect, e.g., by hitting a non-overlapping phase bin of 1/8 of the oscillatory beta cycle.^4^

In the present study, we aimed to address this question in two ways: First, we estimated the circular SD required to hit 2, 4, 8, 16 and 32 equidistant and non-overlapping phase bins of the oscillatory cycle with high precision. Second, we investigated the phase-specific precision of a non-predictive approach on real data, i.e., with EEG-triggered TMS using a novel device based on embedded real-time measurement and stimulation. We hypothesized that this novel online stimulation approach would allow selective phase-targeting across different – also higher – frequency bands with the necessary precision.

## Methods

### Simulation

We estimated the circular SD required to target equidistant and non-overlapping phase bins of the oscillatory cycle with an error rate of less than 1 in 10 000. Specifically, we drew 100 000 draws from a von Mises distribution with varying k. This resulted in 500 simulated datasets (with k from 100 to 1010). To be comparable to other studies on this topic, we calculated the circular SD for each simulated dataset. This resulted in SDs ranging from 0.0005° to 72.8°. We investigated 2, 4, 8, 16 and 32 non-overlapping phase bins. The width of the phase bins was therefore changing with the number of bins. For example, the width was 180° in the case of 2 phase bins and 11.25° in the case of 32 phase bins. Finally, we calculated for each simulated dataset, how often a sample would land outside the width of a phase bin. For each number of phase bin, we also calculated the SD resulting in an error rate of less than 1 in 10 000. We selected this strict threshold to achieve high precision, also when considering large studies with many participants and multiple sessions.

### Real-time measurement and stimulation device

The novel real-time measurement and stimulation device (rtMSD) implemented and investigated here enables us to measure electrophysiological signals, analyze them instantaneously, and return a digital output for triggering external stimulation devices. The device is comprised of separate modules, each with its own defined functionality. The latest version of the rtMSD contains four essential elements in a common housing. The first element is the control chip which ensures real-time communication between all elements. It also drives a set of LEDs to indicate status and functionality of the device and its modules. The second element is a system on a chip (SoC), running a Linux-based operating system. In this environment, custom-written software supports sending data and receiving commands to and from external computers, and performing the signal processing and phase detection in real time. The third element is an analog-to-digital converter which allows the digitization of up to 8 channels of electrophysiological measurements. The last element is a digital input/output connected to two BNC adapters providing TTL input and output and triggering external devices. The first generation of the device used a separate PC instead of the SoC, whereas the current version is completely embedded (see figure 1) and was manufactured in collaboration with neuroConn (Loop-IT, Ilmenau, Germany).

**Figure 1:**
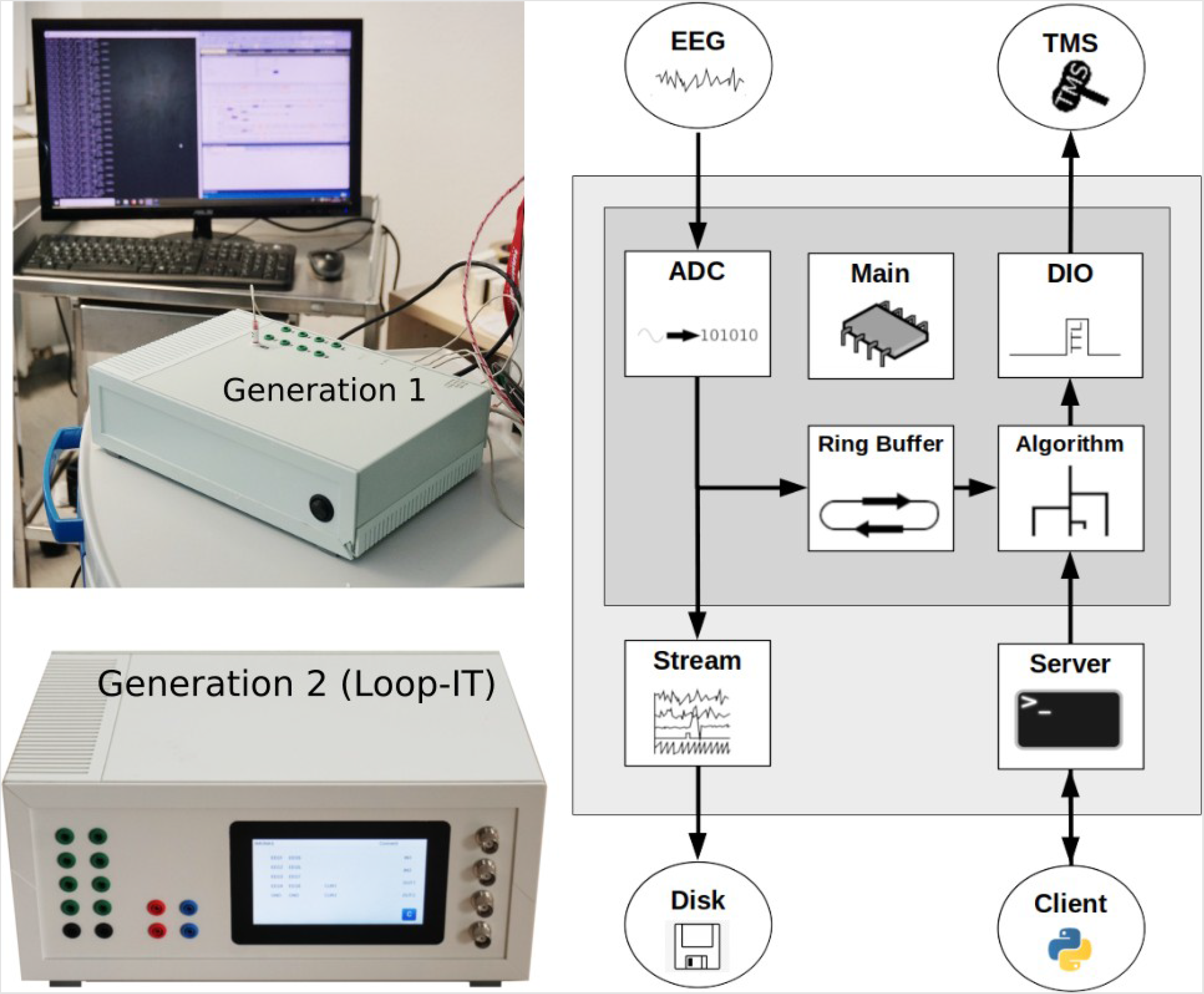
Integrated real-time measurement and stimulation device. The left two images show the first and second generation of the device, and the right diagram shows the internal architecture of both generations. The dark gray box shows the real-time components, the light gray box shows the components with a permissible jitter, and the circles outside the boxes indicate external elements. The EEG is measured, converted and stored in a ringbuffer, which is used by the algorithm to trigger stimulation (e.g., TMS) via a digital input/output interface. The main module realizes real-time performance by cyclic execution and information transfer between the modules. Parameters of the trigger algorithm are shared with a server, which can communicate with an external client for reading and writing the parameters. Raw data and derived measures are streamed online via LabStreamingLayer and can be recorded on an external PC.

### From EEG recording to TMS pulse application

Algorithms for signal processing and analysis as well as phase detection and triggering can be programmed using a high-level language (EN 61131-3 or C/C++) and run on the SoC of the rtMSD directly. The following approach was implemented to evaluate the phase precision of the device. EEG was recorded in a bipolar montage, with two electrodes placed 1 cm anterior and posterior of C3 with the ground electrode on the forehead, and sampled at 1 kHz at a resolution of 24 bit. The bipolar signal was stored in a ring buffer of 500 ms length, implemented as a circular buffer of 500 doubles in memory for improved speed. The window size was defined on the basis of our experience from previous experiments.^1,10^ A discrete Fourier transform for a specific frequency was performed every millisecond with the Görtzel algorithm.^11^ Taking the buffer width of 500ms into account, this provided a frequency resolution of 2 Hz and a phase estimate for each sample. The trigger algorithm considers the current phase and compares it with the phase of the last sample. When the circular arc between the last and the current phase estimate includes the target phase, thereby ascertaining that the target phase has been passed, a trigger signal is delivered. Additionally, the raw EEG data and the calculated phase and trigger channel are streamed together using the LabStreamingLayer. This makes it possible to collect the data on an external PC, where it is then stored on a disk.

### Empirical evaluation

To evaluate the feasibility of this device in practical applications, the 5 V trigger signal was routed via BNC to a TMS device (MagPro X100, MagVenture, Denmark). Since triggering an external device can induce additional latency and inherent jitter, we evaluated the delay of the TMS device by driving it repetitively with a signal generator and measuring the stimulation delay with an oscilloscope. This experiment suggested that the TMS device added ∼65 µs delay, which can be considered negligible for most practical applications.

We evaluated the performance of the rtMSD in 4 right-handed participants (1 female, age M = 23.75, SD = 1.09) who gave their written informed consent prior to participation. The study protocol conformed to the Declaration of Helsinki and was approved by the Ethical Committee of the medical faculty of the University of Tübingen. The study followed the current safety guidelines for application of TMS,^12^ and none of the participants reported side effects.

To study the phase precision of the device over a wide range of frequencies, we investigated 10 different target frequencies (4, 8, 12, 16, 20, 24, 28, 32, 36 and 40 Hz) in randomized order at the target phase of 0°. Stimuli were triggered with an interstimulus interval (ISI) of 3.5-4.5s. We used a jitter of ± 0.5 seconds to reduce anticipation effects. We applied TMS at 120 % resting motor threshold (RMT) to the motor hot spot of the left hemisphere with a pulse width of 280 µs, using biphasic pulses that are considered more effective in inducing MEPs than monophasic pulses.^13^ Due to technical artifacts, 95 trials (11.88 %) were rejected, resulting in overall 705 pulses in the four participants, with 51 to 69 trials per target frequency.

### Performance measures

Bearing in mind that our algorithm triggers only when the target phase has been passed, we expected a linear increase in the phase delay with increasing frequency and compared it to the measured phase delay (see below).

Furthermore, we measured the precision of our stimulation by estimating the circular standard deviation (SD). This entailed cutting 500 ms segments from the recorded signal that ended immediately prior to each trigger and filtering it with a single-order two-way Butterworth filter using a passband of ± 1 Hz around the target frequency. To reduce edge artifacts, the segment was wrap-padded with 1000 samples on each side before filtering. The resulting filtered segments were z-transformed to prevent bias between trials due to unequal signal power. Finally, we averaged all segments per target frequency and calculated the resulting confidence interval at each sample.

### Software Packages

Offline digital processing and statistical analysis were performed using NumPy 1.20.3 and SciPy 1.5.2 on Python 3.7.6 on Linux Mint 20. Additionally, we used pyCircStat 0.0.2 and Statsmodels 0.12.0 and Matplotlib 3.1.3 for visualization.

## Results

The simulation revealed that circular SDs of less than 17.6°, 9.7°, 5.1°, 2.5°, and 1.3° are necessary to hit a specific phase of the oscillatory cycle divided into 2, 4, 8, 16, and 32 non-overlapping phase bins with sufficient precision (i.e., an error rate of smaller than 1/10.000) (fig.2).

**Figure 2:**
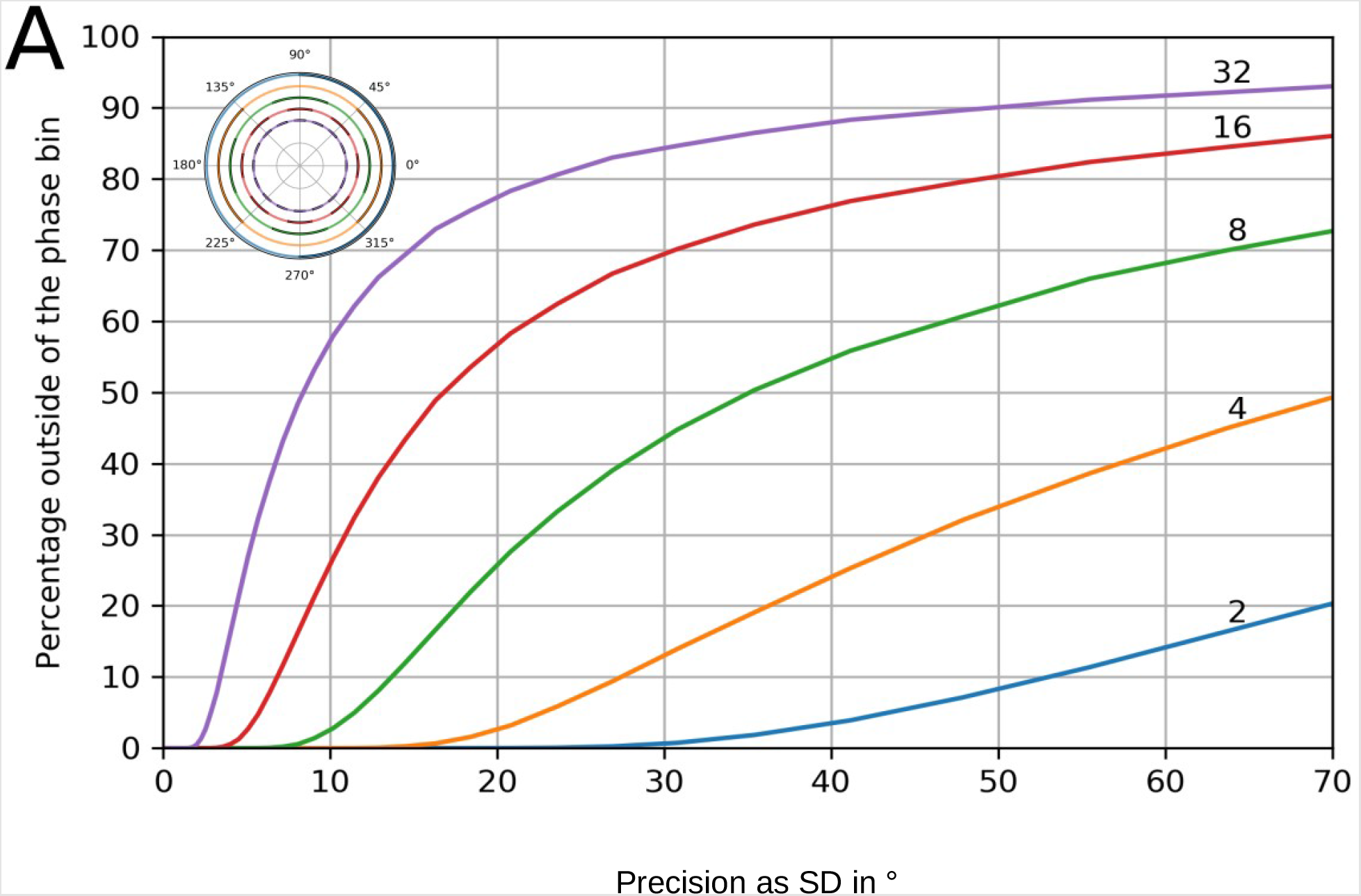
Simulated precision of phase-dependent modulation. The performance curves are plotted with the x-axis showing the circular standard deviance in degrees, indicating the phase precision. The y-axis shows how many samples lie outside of the respective phase bin for a given width on the basis of the number of bins. Each colored line represents a different number of non-overlapping equidistant phase bins, with the bin number ranging from 2 to 32. The inlet in the upper left corner visualizes the number of phase bins for different samplings (from 2 to 32) on the unit circle with dark and light arcs.

The empirical approach completed the measurement, signal-processing and stimulation with a round-trip time of 1 ms, which led to short phase delays for all frequencies between 4 and 40 Hz (figure 3A). Across all frequencies, the spread exhibited a strongly left-skewed distribution (figure 3B). This observation is in line with the expected increase in phase delay as frequency increased, i.e., shorter length of the respective oscillatory cycle. For each frequency investigated, the measured phase was invariably very close to the expected phase delay (see figure 3C), thus allowing for a systemic lag-correction. The grand average of the pre-stimulation segments showed that the bandpass-filtered signal exhibited a sinusoidal modulation, with the stimulation occurring at the peak of the oscillation across frequencies, as expected when triggering at 0° (see figure 3D).

**Figure 3:**
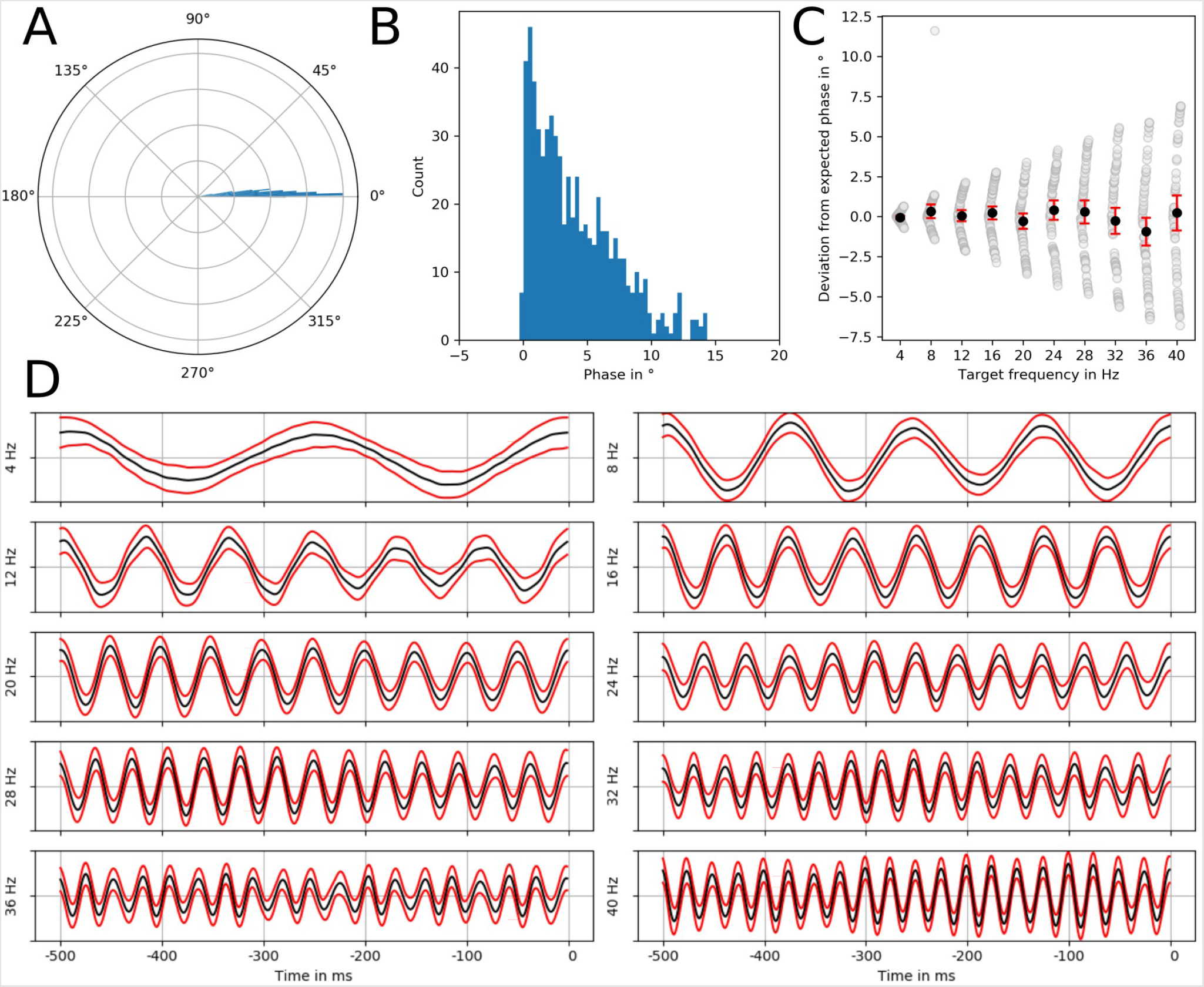
Visualized phase delay and precision (confidence interval) of the integrated real-time measure and stimulation device. The upper row shows the phase delay for the targeted frequencies. (A) The density estimates are given in polar coordinates and arbitrary units (A) and as a count histogram (B). A wheat scatter plot (C) shows the distributed (gray) and average (black) deviance from the expected phase, and the red lines indicating the 95% confidence interval. (D) Band-pass filtered grand average EEG data (black) 500ms before the TMS pulse and the 95% confidence interval (red).

Importantly, our approach achieved very low SDs of 0.4° at 4 Hz to 4.3° at 40 Hz from the targeted phases (table 1). This high precision enabled us to selectively target 32 phases (at 4 Hz), 16 phases (at 8, 12, 16, 20, 24 Hz) and 8 phases (at 28, 32, 36, 40 Hz), respectively, when considering the findings of the simulation.

**Table 1:**
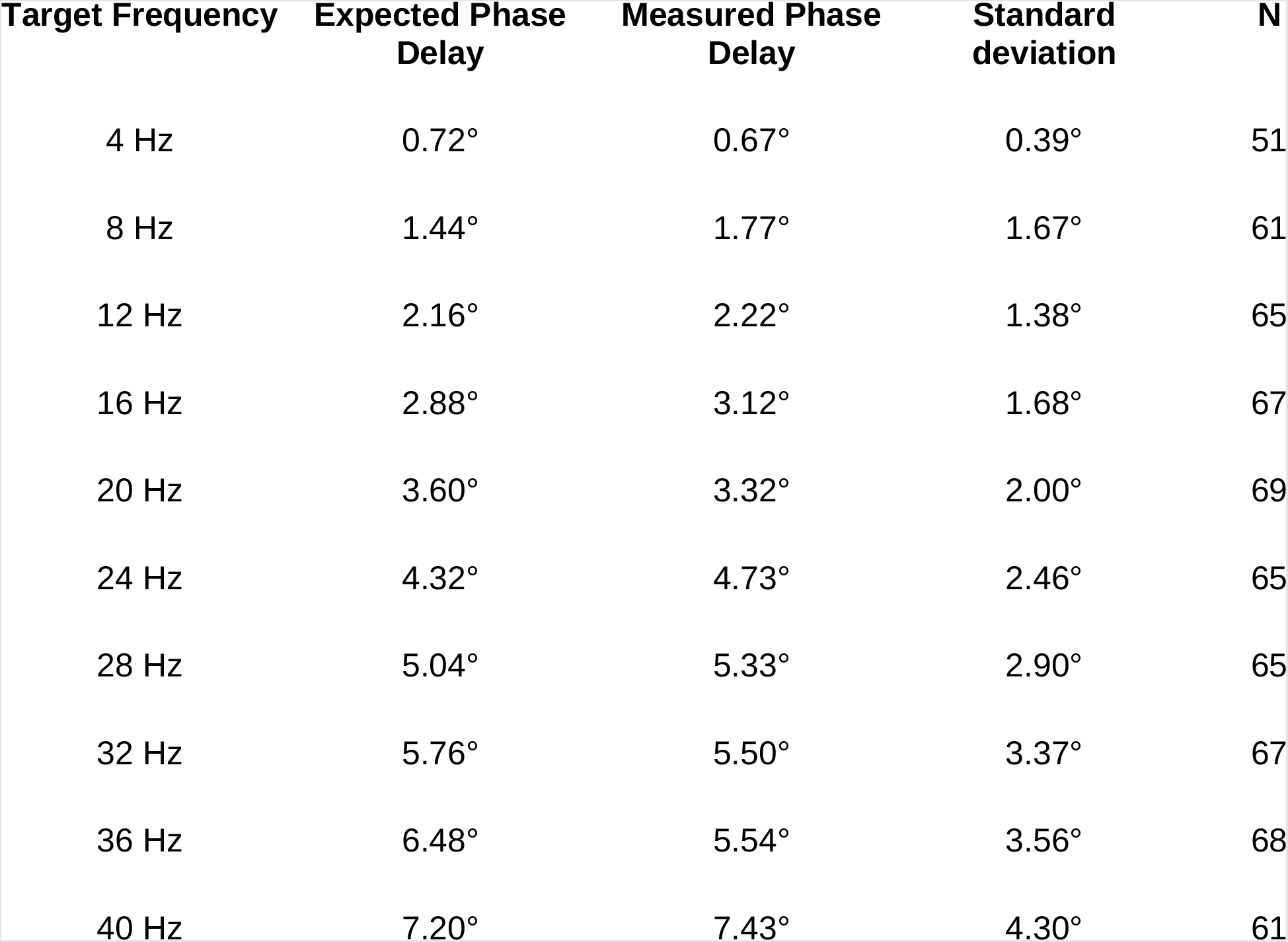
Expected and measured phase delay and standard deviation of the integrated real-time measure and stimulation device for different target frequencies. The overall accuracy consists of the measured phase delay and the standard deviation (precision). With a defined system latency (i.e., measurement, signal-processing and stimulation with a round-time of 1 ms), the expected phase delay increases with increasing target frequency, i.e., shorter length of the respective oscillatory cycle. Since the measured phase delay matched the expected phase delay, a systemic lag-correction of this predictable system delay is possible. The standard deviation (precision), however, is related to the system’s jitter and/or non-stationarity of the target frequency, and is therefore unpredictable. This precision is essential to specifically target the intended phase bin.

| Target Frequency | Expected Phase Delay | Measured Phase Delay | Standard deviation | N |
| --- | --- | --- | --- | --- |
| 4 Hz | 0.72° | 0.67° | 0.39° | 51 |
| 8 Hz | 1.44° | 1.77° | 1.67° | 61 |
| 12 Hz | 2.16° | 2.22° | 1.38° | 65 |
| 16 Hz | 2.88° | 3.12° | 1.68° | 67 |
| 20 Hz | 3.60° | 3.32° | 2.00° | 69 |
| 24 Hz | 4.32° | 4.73° | 2.46° | 65 |
| 28 Hz | 5.04° | 5.33° | 2.90° | 65 |
| 32 Hz | 5.76° | 5.50° | 3.37° | 67 |
| 36 Hz | 6.48° | 5.54° | 3.56° | 68 |
| 40 Hz | 7.20° | 7.43° | 4.30° | 61 |

## Discussion

We investigated the phase-specific precision of state-dependent neuromodulation on the basis of integrated real-time measurement and brain stimulation. By completing measurement, signal-processing and stimulation with a round-time of 1 ms, this novel non-predictive approach achieved high empirical precision in the real-life scenario of EEG-triggered TMS. Specifically, the standard deviations across ten different frequencies were 0.4° at 4 Hz to 4.3° at 40 Hz. According to our simulations, this precision would facilitate the selective targeting of 8 distinct phases in all investigated frequencies up to 40 Hz. This precision would be sufficient to achieve the necessary phase specificity of TMS to maximize MEP increases in accordance with previous post-hoc estimations.^4^

By contrast, previous autoregressive (AR) model-based approaches using pre-stimulus estimation of the brain state for EEG-triggered TMS do not achieve such high precision as they require the application of prediction algorithms to compensate for latencies between measurement and stimulation. Accordingly, previous studies applied EEG-triggered TMS to investigate research questions that could be addressed with less temporal precision; e.g., by applying EEG-triggered TMS on the basis of (a) high and low oscillatory power levels in the beta band (16-22 Hz) and (b) positive and negative peaks of the slow (< 1Hz) or alpha (8-12 Hz) oscillatory cycle to study instantaneous and lasting MEP amplitude changes: Specifically, brain state-dependent TMS, when controlled by volitionally modulated low sensorimotor beta power levels, induced a robust MEP amplitude increase.^1^ By contrast, when the very same stimulation pattern was applied independent of the brain state, a decrease in corticospinal excitability ensued. When TMS was applied during sleep, targeting depolarized vs. hyperpolarized phases of slow oscillations was associated with an increase in instantaneous MEP amplitudes.^14^ During wakefulness, sensorimotor alpha oscillations at rest were used to trigger TMS, targeting the negative vs. positive peak of the oscillatory cycle and thus leading to a lasting increase vs. no change in MEP amplitude in a preselected group of participants with high sensorimotor alpha power.^2^ However, this finding was not replicated when the same approach was applied in non-selected individuals.^3^ Importantly, these online approaches hit the targeted negative/positive peaks of the alpha cycle with standard deviations of 55°/53°^2^ and 48°/52°.^3^

According to the simulation of this study (figure 2), a considerable number of stimuli would be outside the target phase bin when imprecise stimulation with a SD of ∼50° is applied with previous approaches. Specifically, >30% of stimuli would be off-target for a phase resolution of 4 bins, which is necessary to estimate sinusoidal modulation. Furthermore, >60% of stimuli would be off-target for a phase resolution of 8 bins, which has been shown to be necessary to capture phase-dependent effects of corticospinal excitability in the oscillatory beta-band.^4^

The inconsistencies of these previous observations may therefore be related to different factors: On the one hand, the relative imprecision of hitting the targeted phase of the alpha cycle may have prevented more robust and reproducible findings with regard to TMS induced plastic changes. On the other hand, the intrinsic targeting error of current approaches is amplified when higher frequency bands are investigated, and thus limits studying the phase-dependency of frequencies that may shape the timing of voluntary movements and determined corticospinal excitability in earlier offline analyses.^15-18^

Moreover, the interaction between oscillatory phase and power may influence corticospinal excitability to an extent unexplained by phase or power alone.^19^ Also, resolving current contradictions of phase dependency in different frequencies^15,20^ may necessitate high-resolution sampling along the oscillatory cycle (e.g., by comparing 8 equidistant phases in each investigated frequency) to allow for robust modeling of input-output relationships.^21^

Furthermore, minor differences in the methodological choices may critically affect the sensitivity to detect the complex relationship between oscillatory activity and corticospinal excitability^22^ and even lead to erroneous phase estimations.^23^

### Limitations and perspectives

The technological feasibility of highly precise phase-specific stimulation does not per se lead to the intended stimulation effects and desired outcome. Before translating this approach to broad scientific and clinical application, relevant open questions need to be addressed: Although hitting the intended phase precisely may result in reduced variability of instantaneous stimulation effects, as demonstrated in previous post-hoc analyses;^17^ these less variable stimulation effects do not necessarily lead to cumulative stimulation effects and plastic changes after repetitive application. Moreover, it needs to be clarified how consistent phase-dependent stimulation effects are in cross-validation experiments, e.g., with out-of-sample evaluation within one session, across different sessions and days, and between individuals with regard to the optimal frequency and phase bin of the oscillatory cycle. ^21^ Furthermore, stimulation pulses may also be timed to the most sensitive phase with novel computation methods that are more precise and faster than previous prediction approaches by modeling a linear oscillator and recomputing the phase of this virtual oscillator into the analyzed signal.^24,25^ Notably, the sinusoidal nature of cortical oscillations, and whether they might be better characterized by their non-sinusoidality, is currently a matter of some debate.^26,27^ However, such open questions may now be addressed with this novel non-predictive stimulation approach.

In conclusion, integrated real-time measurement and brain stimulation circumvented the need for prediction and allowed state-informed stimulation with high precision. The applied approach of EEG-triggered TMS resulted in selective targeting of 8 distinct phases in the investigated frequencies of up to 40 Hz, thereby attaining the necessary specificity to maximize instantaneous stimulation effects. Future studies need to clarify whether this will also lead to increased cumulative stimulation effects and plastic changes, and whether phase-specific stimulation may influence brain disorders that are characterized by aberrant neural oscillations.^28-31^

## Authorship contribution statement

Robert Guggenberger: Conceptualization, Methodology, Software, Data curation, Formal analysis, Writing - original draft, Visualization. Julian-Samuel Gebühr: Methodology, Software, Investigation, Data curation, Writing - review & editing. Marius Keute: Methodology, Software, Writing - review & editing. Alireza Gharabaghi: Conceptualization, Writing - original draft, Supervision, Project administration, Funding acquisition.

## Declaration of Competing Interest

The authors declare no conflict of interests.

## Acknowledgments

This work was supported by the German Federal Ministry of Education and Research [BMBF16SV8174, INERLINC]. We acknowledge support from the Open Access Publishing Fund of the University of Tuebingen.

## Data availability statement

The data that support the findings of this study are available for researchers from the first author upon reasonable request.

## Notes

### Competing Interest Statement

The authors have declared no competing interest.

## References

1. Kraus D, Naros G, Bauer R, Khademi F, Leão MT, Ziemann U, Gharabaghi A. (2016) Brain State-Dependent Transcranial Magnetic Closed-Loop Stimulation Controlled by Sensorimotor Desynchronization Induces Robust Increase of Corticospinal Excitability. Brain Stimul. 9:15–424.

2. Zrenner C, Desideri D, Belardinelli P, Ziemann U (2018) Real-time EEG-defined excitability states determine efficacy of TMS-induced plasticity in human motor cortex. Brain Stimul 11: 374–389.

3. Madsen KH, Karabanov AN, Krohne LG, Safeldt MG, Tomasevic L, Siebner HR (2019) No trace of phase: Corticomotor excitability is not tuned by phase of pericentral mu-rhythm. Brain Stimul 12:1261–1270.

4. Khademi F, Royter V, Gharabaghi A (2019) State-dependent brain stimulation: Power or phase? Brain Stimul 12:296–299.

5. Chen LL, Madhavan R, Rapoport BI, Anderson WS (2013) Real-time brain oscillation detection and phase-locked stimulation using autoregressive spectral estimation and time-series forward prediction. IEEE Trans Biomed Eng. 60:753–62.

6. Onojima T, Kitajo K. (2021) A state-informed stimulation approach with real-time estimation of the instantaneous phase of neural oscillations by a Kalman filter. J Neural Eng. 9;18(6).

7. Shakeel A, Onojima T, Tanaka T, Kitajo K (2021) Real-Time Implementation of EEG Oscillatory Phase-Informed Visual Stimulation Using a Least Mean Square-Based AR Model. J Pers Med. 11;11(1):38

8. Shirinpour, S., Alekseichuk, I., Mantell, K., & Opitz, A. (2020). Experimental evaluation of methods for real-time EEG phase-specific transcranial magnetic stimulation. Journal of neural engineering, 17(4), 046002.

9. McIntosh JR, Sajda P (2020) Estimation of phase in EEG rhythms for real-time applications. J Neural Eng. 2020 Jun 2;17(3):034002.

10. Raco V, Bauer R, Tharsan S, Gharabaghi A (2016) Combining TMS and tACS for Closed-Loop Phase-Dependent Modulation of Corticospinal Excitability: A Feasibility Study. Front Cell Neurosci 10:143.

11. Sorensen HV, Burrus CS, Jones DL (1988) A new efficient algorithm for computing a few DFT points. In: 1988., IEEE International Symposium on Circuits and Systems, pp 1915–1918.

12. Rossi S, Antal A, Bestmann S, Bikson M, et al. (2021) Safety and recommendations for TMS use in healthy subjects and patient populations, with updates on training, ethical and regulatory issues: Expert Guidelines. Clin Neurophysiol. 132:269–306.

13. Terao Y, Ugawa Y (2002) Basic Mechanisms of TMS: J Clin Neurophysiol 19:322–343.

14. Bergmann TO, Mölle M, Schmidt MA, Lindner C, Marshall L, Born J, Siebner HR (2012) EEG-guided transcranial magnetic stimulation reveals rapid shifts in motor cortical excitability during the human sleep slow oscillation. J Neurosci 32:243–253.

15. Khademi F, Royter V, Gharabaghi A (2018) Distinct Beta-band Oscillatory Circuits Underlie Corticospinal Gain Modulation. Cereb Cortex 28:1502–1515.

16. Naros G, Lehnertz T, Leão MT, Ziemann U, Gharabaghi A (2020) Brain State-dependent Gain Modulation of Corticospinal Output in the Active Motor System. Cereb Cortex. 10; 30:371–381

17. Torrecillos, F., Falato, E., Pogosyan, A., West, T., Lazzaro, V. D., & Brown, P (2020). Motor Cortex Inputs at the Optimum Phase of Beta Cortical Oscillations Undergo More Rapid and Less Variable Corticospinal Propagation. J Neurosci 40, 369–381.

18. Hussain SJ, Vollmer MK, Iturrate I, Quentin R. (2022) Voluntary Motor Command Release Coincides with Restricted Sensorimotor Beta Rhythm Phases. J Neurosci. 20; 42:5771–5781.

19. Hussain, S. J., Claudino, L., Bönstrup, M., Norato, G., Cruciani, G., Thompson, R., … & Cohen, L. G. (2019). Sensorimotor oscillatory phase–power interaction gates resting human corticospinal output. Cerebral Cortex, 29(9), 3766–3777.

20. Wischnewski M, Haigh ZJ, Shirinpour S, Alekseichuk I, Opitz A. (2022) The phase of sensorimotor mu and beta oscillations has the opposite effect on corticospinal excitability. Brain Stimul. 15:1093–1100.

21. Keute M, Gebühr JS, Guggenberger R, Trunk BH, Gharabaghi A (2022) Phase-specific stimulation reveals consistent sinusoidal modulation of human corticospinal excitability along the oscillatory beta cycle. (under review)

22. Karabanov AN, Madsen KH, Krohne LG, Siebner HR. (2021) Does pericentral mu-rhythm “power” corticomotor excitability? A matter of EEG perspective. Brain Stimul. 14:713–722.

23. Khademi F, Royter V, Ziegler L, Gharabaghi A (2022) Resolving equivocal gain modulation of corticospinal excitability. (under review)

24. Rosenblum, M. (2020). Controlling collective synchrony in oscillatory ensembles by precisely timed pulses. Chaos: An Interdisciplinary Journal of Nonlinear Science, 30(9), 093131.

25. Busch JL, Feldmann LK, Kühn AA, Rosenblum M. (2022) Real-time phase and amplitude estimation of neurophysiological signals exploiting a non-resonant oscillator. Exp Neurol. 347:113869.

26. Cole SR, Voytek B (2017) Brain Oscillations and the Importance of Waveform Shape. Trends Cogn Sci 21:137–149.

27. Schaworonkow N, Nikulin VV (2019) Spatial neuronal synchronization and the waveform of oscillations: Implications for EEG and MEG Battaglia FP, ed. PLOS Comput Biol 15: e10070551:

28. Azodi-Avval, R., & Gharabaghi, A. (2015). Phase-dependent modulation as a novel approach for therapeutic brain stimulation. Frontiers in computational neuroscience, 9, 26.

29. Cagnan H, Pedrosa D, Little S, Pogosyan A, … & Brown P. (2017) Stimulating at the right time: phase-specific deep brain stimulation. Brain. 140:132–145.

30. Holt AB, Kormann E, Gulberti A, Pötter-Nerger M, … & Sharott A. (2019) Phase-Dependent Suppression of Beta Oscillations in Parkinson’s Disease Patients. J Neurosci. 6; 39:1119–1134.

31. McNamara, C. G., Rothwell, M., & Sharott, A. (2020). Phase-dependent closed-loop modulation of neural oscillations in vivo. BioRXiv.

